# Trim7 does not have a role in the restriction of murine norovirus infection in vivo

**DOI:** 10.1101/2024.10.17.618898

**Authors:** Mridula Annaswamy Srinivas, Linley R. Pierce, Mikayla C. Olson, Shelly J. Roberston, Gail L. Sturdevant, Sonja M. Best, Robert C. Orchard

## Abstract

Trim7 is an E3 ubiquitin ligase that was recently identified as a central regulator of host- viral interactions with both pro-viral and anti-viral activity in cell culture. As an inhibitor, Trim7 overexpression ubiquitinates viral proteins by recognizing C-terminal glutamines that are hallmarks of 3C-like protease cleavage events. Here we sought to determine the physiological impact of Trim7 in resolving murine norovirus (MNV) infection of mice as MNV is potently inhibited by Trim7 in vitro. Utilizing two independently derived Trim7 deficient mouse lines we found no changes in the viral burden or tissue distribution of MNV in both an acute and persistent model of infection. Additionally, no changes in cytokine responses were observed after acute MNV infection of Trim7-deficient mice. Furthermore, removal of potentially confounding innate immune responses such as STING and STAT1 did not reveal any role for Trim7 in regulating MNV replication. Taken together, our data fails to find a physiological role for Trim7 in regulating MNV infection outcomes in mice and serves as a caution for defining Trim7 as a broad acting antiviral.

**Importance:** Intrinsic antiviral molecules that restrict viral replication are important drivers of viral evolution and viral tropism. Recently, Trim7 was shown to provide cell intrinsic protection against RNA viruses, including murine norovirus. Biochemically, Trim7 recognizes the cleavage product of viral proteases, suggesting a novel and broad mechanism to restrict viral replication. Here, we tested whether Trim7 had a physiological role in restricting murine norovirus replication in mice. Unexpectedly, we found no impact of viral replication or innate immune responses during murine norovirus infection. Our findings urge caution in defining Trim7 as a broad antiviral factor in the absence of in vivo evidence.

## Introduction

Noroviruses are non-enveloped positive-sense single stranded RNA viruses that are a leading cause of infectious gastroenteritis worldwide^1,2^. Norovirus infections in humans are typically self-resolving but cause a significant economic burden^3^. Children and immunocompromised individuals can be affected by more severe, recurrent and persistent infections^4,5^. Despite their high infectivity and burden, there is currently no vaccine or treatment for noroviral gastroenteritis due to difficulties in culturing human noroviruses (HNoV). Stem-cell derived enteroid cultures enable growth of HNoV in vitro and hold promise to accelerate therapeutic discoveries^6–9^. However, there are no accessible small animal models to study HNoV infections in vivo^10^. Murine norovirus (MNV) has emerged as a model system to study HNoV due to its ability to infect mice and replicate in standard cell culture conditions^11–13^. The MNV model system is a robust platform to study host-virus interactions and immune factors against noroviruses in a natural setting.

Previously, we identified a host protein Tripartite motif-containing protein 7 (Trim7) as a strong antiviral protein against MNV replication^14^. Trim7 is an E3 ubiquitin ligase that was originally described as an interactor of glycogenin^15^. Trim7 preferentially binds and ubiquitinates substrates with a C-terminal glutamine residue^16–18^. In recent years, Trim7 has been widely studied for its role in viral replication and host defense systems with reports of both proviral and antiviral activities. For example, Trim7 ubiquitinates Zika virus envelope protein E, enhancing viral entry into host cells^19^. Others report that Trim7 ubiquitinates stimulator of interferon genes (STING) and mitochondrial antiviral signaling protein (MAVS) leading to a reduced innate immune response and decreased protection against infection^20,21^. Contrastingly, Trim7 has antiviral activity towards enteroviruses via ubiquitination of non-structural protein 2BC of coxsackievirus CVB3^22^. The CVB3 3C protease cleaves Trim7 to antagonize its antiviral function^23^. With respect to noroviruses, we and others have further demonstrated by infection and biochemical studies that Trim7 can target norovirus non-structural proteins NS6 and NS3^17,18,24^. 3C-like viral proteases preferentially cleave substrates at a glutamine residue^25^, placing a potential for Trim7 to be a broad regulator of 3C-protease cleavage products. However, the antiviral role of Trim7 in the physiological setting has not been tested. Given the complex relationship between pro- and anti-viral facets of Trim7 biology, it is important to define how this protein at the nexus of intrinsic immunity impacts viral infection in vivo.

MNV strains have disparate infection outcomes in mice that have been used to model different aspects of HNoV infections^11,12^. Certain strains of MNV such as MNV^CW3^ cause an acute infection of both intestinal and extraintestinal tissues via infection of immune cells^26,27^. These infections are typically self-resolving but can persist in the absence of the adaptive immune system and are lethal in interferon deficient settings, such as STAT1^-/-^ mice^12,28^. Acute infections by MNV have been used to model the acute phase of HNoV infections. In contrast, some strains of MNV establish a persistent, enteric infection that is confined to the gastrointestinal tract^29^. The prototypical persistent strain MNV^CR6^ infects tuft cells and evades both innate and adaptive immunity through mechanisms that are still not well understood^30–36^. Persistent MNV infections mirror the long-term, asymptomatic shedding observed for HNoV^37^. The diversity of MNV phenotypes provides not only an opportunity to explore different properties of norovirus pathogenesis but also the role of host factors such as Trim7 in restricting MNV in a cell or tissue specific manner.

To determine the physiological impact of Trim7 in resolving both acute and persistent norovirus infections in mice, we utilized two independently derived Trim7 deficient mouse lines. Despite robust restriction of MNV in vitro by overexpressed Trim7, we fail to observe any change in MNV replication in Trim7 deficient mice. Furthermore, no changes in cytokine production were observed during MNV infection when Trim7 was absent. Elimination of potentially confounding immune pathways such as STAT1 or STING did not reveal any physiological role for Trim7 in restricting MNV replication. While Trim7 may play an important role in a yet to be tested MNV infection system, caution is urged in defining Trim7 as a broad acting antiviral recognizing the products of 3C-like protease cleavage events.

## Results

### Generation of Trim7 knockout mouse lines

To assess the role of Trim7 in vivo, we acquired two independent Trim7 deficient mouse lines to circumvent the challenges in detecting endogenous Trim7 protein expression. The first line (BL6/Trim^+1/+1^) has a single nucleotide insertion in exon 1 leading to a premature stop codon and has been previously characterized^19^ (**Figure 1A**). For added rigor, we generated a second Trim7 mouse with a 12kb deletion spanning exons 1 through 7 of Trim7 (BL6ΔTrim7; **Figure 1A**). We have not been able to robustly and reproducibly detect endogenous Trim7 via antibodies from multiple manufacturers in any of our mouse lines or tissues (data not shown). **Figure 1B** is a representative example with one of these antibodies demonstrating a failure to detect a specific Trim7 band. Thus, we designed a qRT-PCR assay targeting exon 1 of Trim7 to determine the tissue distribution of Trim7 in wild-type mice (**Figure 1A**). Consistent with previous results^19,22^, we detect high levels of Trim7 in the heart, leg muscle, and kidney (**Figure 1C)**. We then examined the expression level of Trim7 at sites where MNV replicates. Our data demonstrates low, but detectable levels of Trim7 in the ileum and colon while higher levels at extraintestinal sites like the spleen, liver, and lung (**Figure 1C**). Using this qRT-PCR assay we also confirmed the loss of RNA expression of Trim7 in the BL6ΔTrim7 mice (**Figure 1D**).

**Figure 1.**
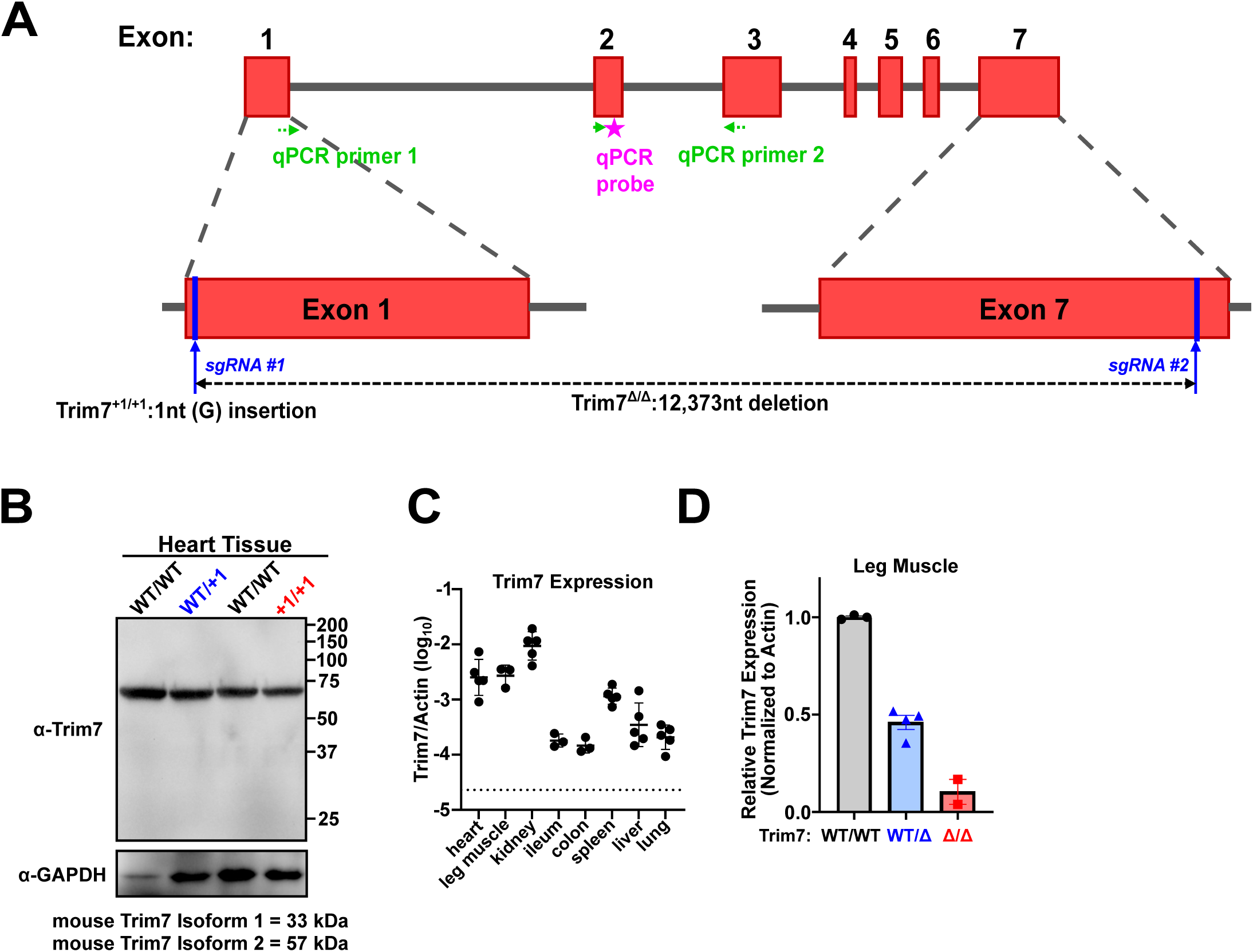
Overview of Trim7 deficient lines used in this study **A)** Cartoon of the mouse Trim7 locus. sgRNAs 1 and 2 were used to generate two Trim7 deficient mouse lines. Trim7^+1/+1^ has a one nucleotide insertion in exon 1 and has been previously reported^19^. Trim7Δ/Δ has a 12,373 nucleotide deletion between exons 1 and 7. Also marked are the qPCR primers and probe used for validation. qPCR primer 1 spans the exon 1/2 junction. **B)** Representative western blot of Trim7 of heart tissue from the indicated mouse lines demonstrating a lack of specific band at the predicted molecular weight of Trim7 isoforms. **C)** Trim7 expression from indicated tissues from wild-type BL6 mice as measured by qPCR and normalized by concurrent quantification of actin transcripts. Dotted line represents the limit of detection and each dot represents an individual mouse. **D)** Trim7 expression in leg muscle relative to wild-type mice after normalization of actin values. Each dot represents an individual mouse.

### No detectable role for Trim7 in restriction of acute murine norovirus infection in vivo

We first tested whether endogenous Trim7 restricts acute, systematic norovirus infection in vivo. We inoculated WT BL6 and BL6/Trim7^+1/+1^ littermates with MNV^CW3^ and harvested tissues 7 days post-infection. Consistent with previous results, we find MNV genomes in mesenteric lymph nodes (MLN), spleen, and liver in wild-type mice (**Figure 2A**). MNV^CW3^ does not infect the colon and poorly infects the ileum. Deficiency of Trim7 in the infected mice does not alter the sites of infection or the burden of MNV in these mice (**Figure 2A**). We also found similar results comparing BL6ΔTrim7 heterozygous and knockout littermates infected with MNV^CW3^ (**Figure 2B**). Overall, these data fail to demonstrate an impact of Trim7 on acute MNV replication in vivo.

**Figure 2.**
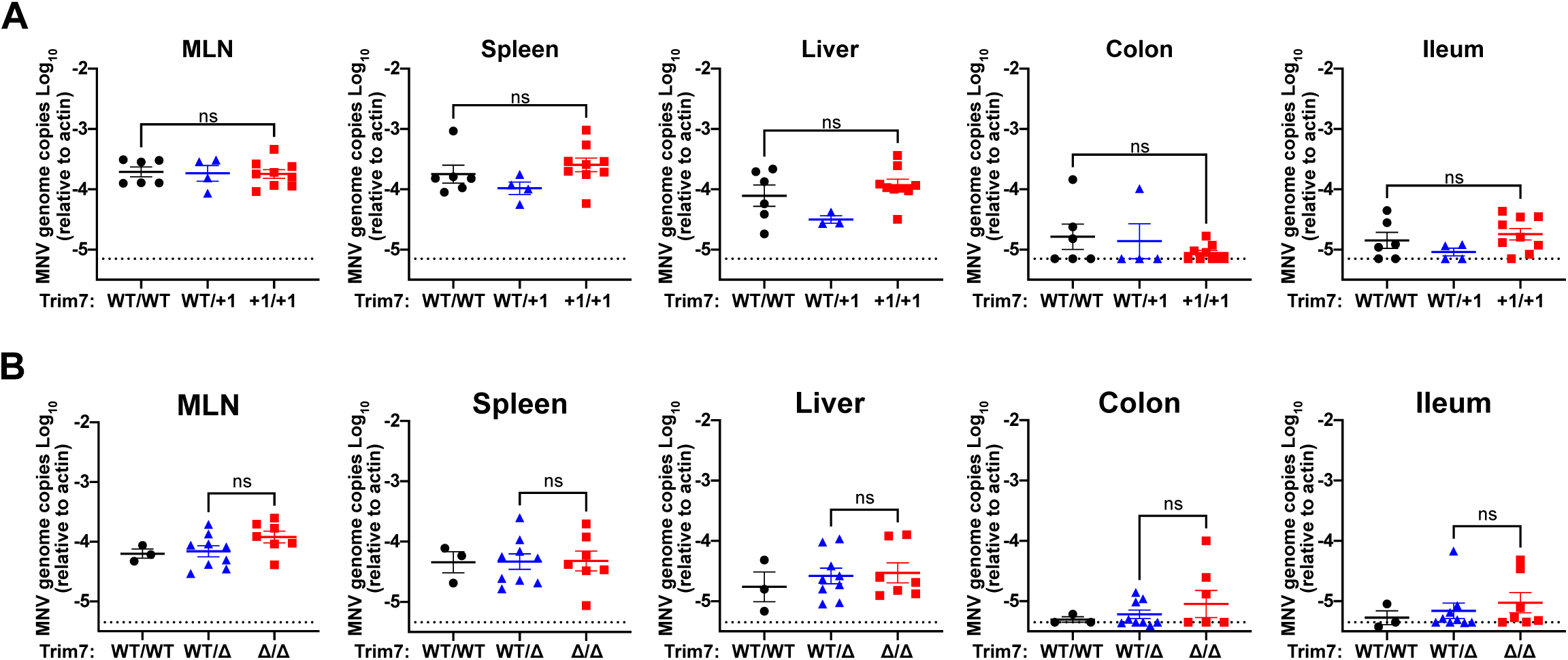
Trim7 does not impact the replication or spread of acute murine norovirus infection *in vivo*. C57BL/6-Trim7^+1/+1^ mice **(A)** or C57BL/6-Trim7^Δ/Δ^ mice (**B**) and respective littermate controls were inoculated with 5 × 10^6^ PFU of MNV^CW3^ and euthanized 7 days post- infection. Tissue titers for mesenteric lymph nodes (MLN), spleen, liver, colon, and ileum were analyzed via qPCR for MNV genome copies and normalized to actin.

### No impact of Trim7 on MNV persistence in gastro-intestinal tissues

Persistent strains of MNV have a distinct tropism of tuft cells thus Trim7 may have an impact on persistent strains rather than acute strains^29,33,34^. To evaluate the impact of Trim7 on the gastro-intestinal tissue infection and persistence of MNV, we infected WT and BL6/Trim7^+1/+1^ littermate-matched mice with MNV^CR6^ and monitored viral load over 21 days of infection. We confirmed successful infection, replication, and persistence of MNV infection in these mice by determining viral load in the feces from day 3 to day 21 post-infection (**Figure 3A**). Trim7 deficiency had no impact on fecal shedding at either early or persistent time points (**Figure 3A**). The colon and MLN harbor high levels of MNV^CR6^ and Trim7 deficiency did not alter the amount of virus detected at these sites (**Figure 3B and 3C**). MNV^CR6^ does not robustly infect the ileum, spleen, or liver of immunocompetent animals^29^. BL6/Trim7^+1/+1^ animals had similarly low or undetectable amounts of MNV genomes in these tissues as compared to their Trim7 sufficient littermate controls (**Figure 3D-3F)**. Taken together these data suggest that Trim7 has no impact on gastrointestinal infection and persistent by MNV^CR6^, nor does Trim7 deficiency enable a new tissue niche at persistent time points.

**Figure 3.**
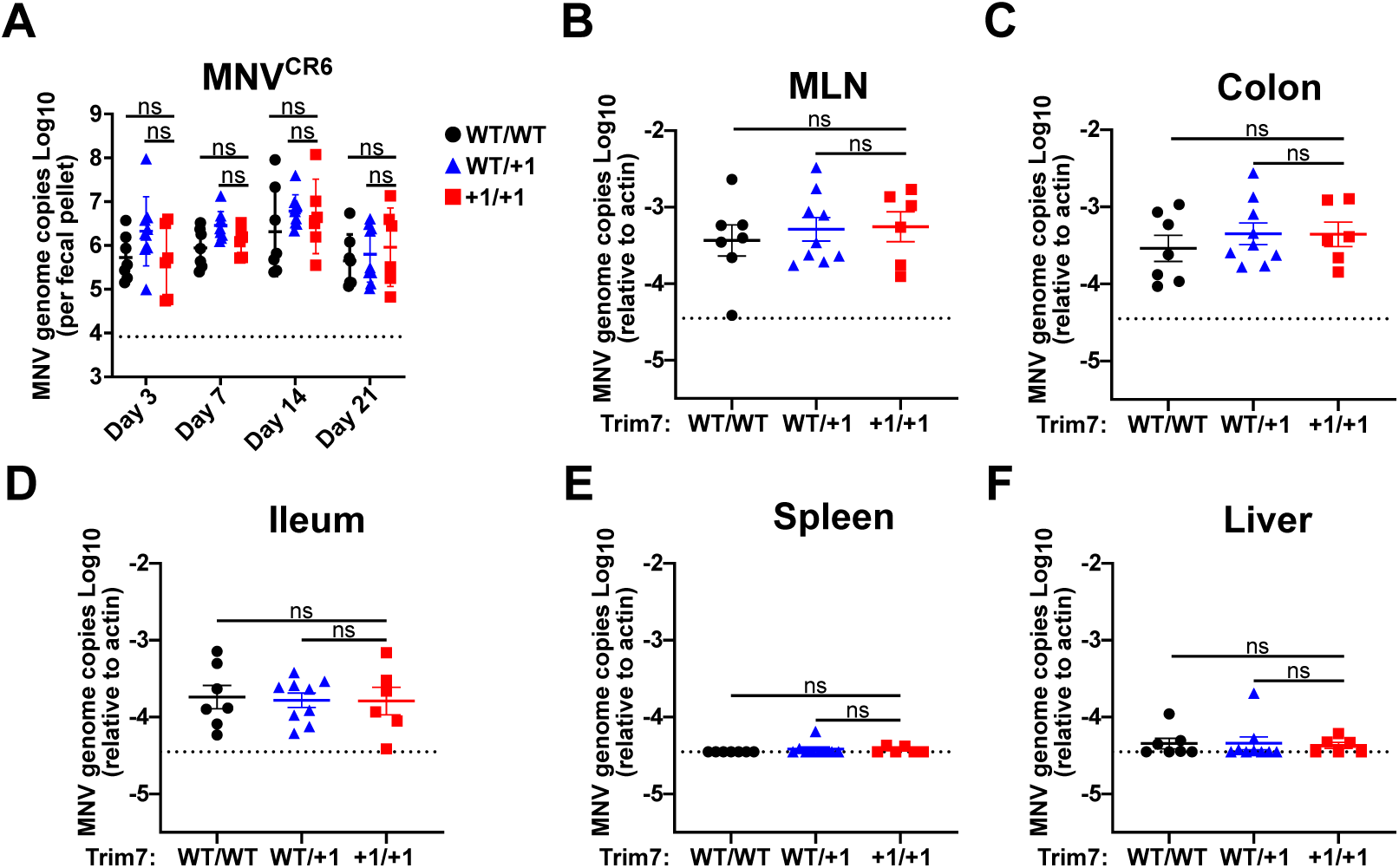
Trim7 does not impact the replication or spread of persistent enteric MNV C57BL/6-Trim7^+1/+1^ mice and littermate controls were inoculated with 1 × 10^6^ PFU of MNV^CR6^ and MNV genome copies were enumerated from fecal samples 3, 7, 14, or 21 days-post infection via qPCR **(A)**. 21 days post-infection, animals were sacrificed and MNV burden assed by measuring the genome copies in the MLN (**B**), colon (**C**), ileum (**D**), spleen (**E**), and liver (**F**) via qPCR. All samples were normalized relative to actin. Data are shown as mean ± S.D. from three independent experiments with 3-9 mice per group. ns, ** *P* < 0.01, *** *P* < 0.001, **** *P* < 0.0001, Mann-Whitney’s test.

### Trim7 does not impact innate immune response to acute MNV infection

Previous studies have shown that Trim7 inhibits innate immune responses by regulating the expression of innate immune sensors STING and MAVS^20,21^. In these studies, Trim7 deficient mice infected with RNA or DNA viruses produced more type I interferons and other pro-inflammatory cytokines and chemokines like TNF-a, IFN-b, Cxcl10 and ISG56^20,21^. MNV infection also leads to interferon stimulation and upregulation of cytokine/chemokine responses^12,41,42^. We investigated the innate immune response to MNV^CW3^ infection in MLN, spleen and liver of BL6/Trim7^+1/+1^ mice at 7 days post-infection by measuring the induction of cytokines via qPCR. No significant differences were observed in the transcript levels of IFN-b1, IL-6, Ifit1 or Cxcl10 in any of the tested tissues between infected BL6/Trim7^WT/WT^ and BL6/Trim7^+1/+1^ mice (**Figure 4A-4D**). These data indicate that Trim7 does not impact the innate immune response to acute MNV infection.

**Figure 4.**
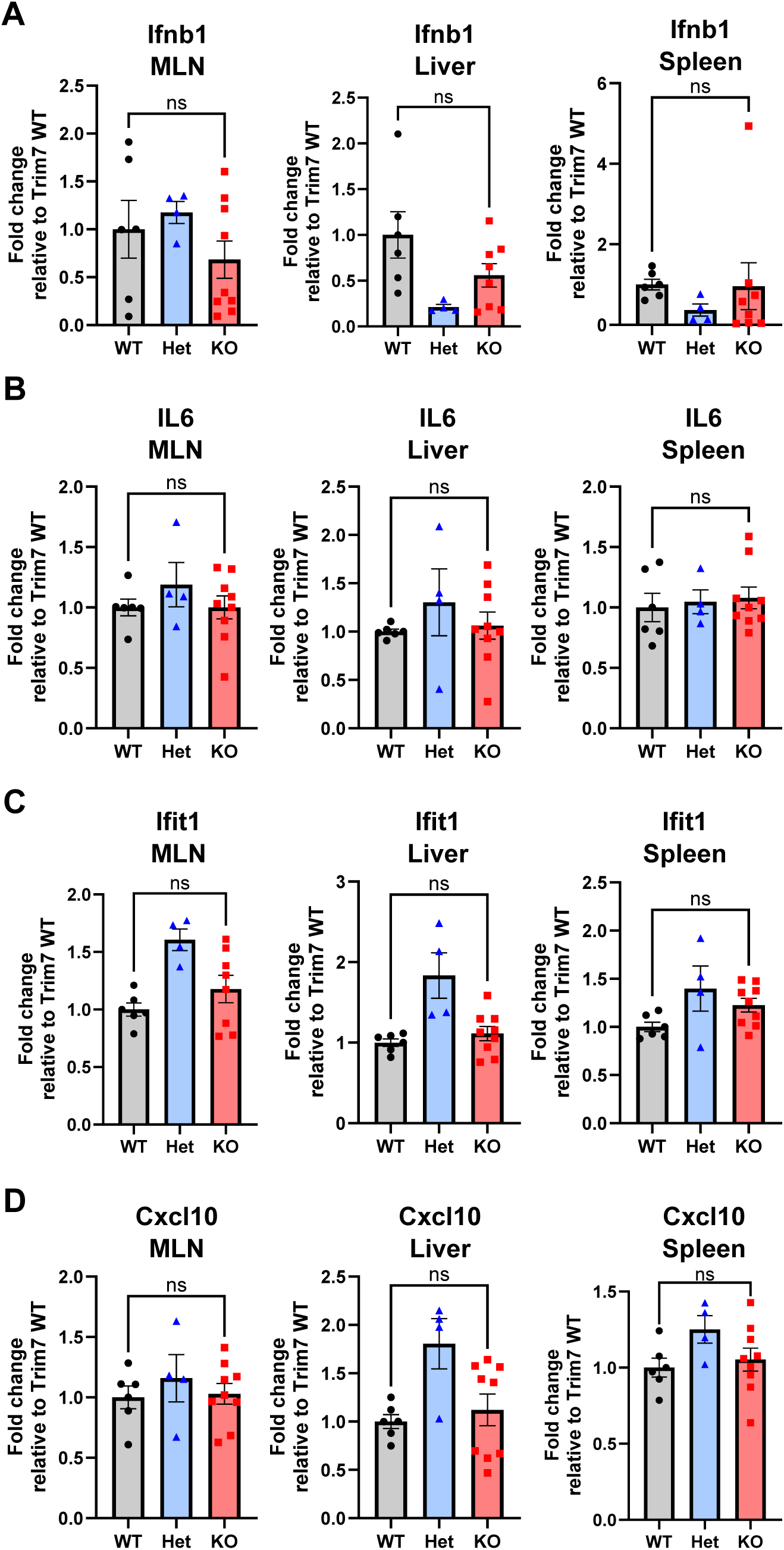
Trim7 does not affect innate immune response to MNV^CW3^ infection. C57BL/6-Trim7^+1/+1^ mice and littermate controls were inoculated with 5 × 10^6^ PFU of MNV^CW3^ and euthanized 7 days post-infection and indicated tissues were collected. **(A)** Ifnb1, **(B)** IL6, **(C)** Ifit1 and **(D)** Cxcl10 transcript copies were determined via qPCR and normalized to actin levels. The data are plotted relative to the average of the quantities in WT mice post-MNV infection. Data are shown as mean ± S.D. from three independent experiments with 4-9 mice per group. ns, ** *P* < 0.01, *** *P* < 0.001, **** *P* < 0.0001, one- way ANOVA with Tukey’s multiple comparison test.

### STING and STAT1 dependent innate pathways do not mask a role of Trim7 in restricting acute MNV infection

MNV can activate the cGAS/STING pathway by inducing the release of mitochondrial DNA from infected cells which in turn can restrict MNV replication in vitro. Whether this pathway is relevant to MNV infection in mice has not been evaluated^43,44^. Trim7 has been reported to target STING for ubiquitination and degradation, thus serving as a proviral factor for viruses^21^. Consequently, it is possible that the physiological effect of Trim7 is masked by the greater impact of Trim7 regulation of STING-mediated restriction of MNV. To eliminate this possible confounding pathway, we crossed the BL6/Trim7^+1/+1^ mice onto a STING^Gt/Gt^ background. In doing so we generated Trim7 sufficient and deficient animals both in the absence of STING. We orally inoculated STING^Gt/Gt^ Trim7^+1/+1^ or littermate STING^Gt/Gt^ Trim7^WT/+1^ controls with MNV^CW3^ and assessed replication in the MLN, spleen, liver, colon, or ileum via qPCR 7 days after infection. In each tissue, MNV genomes were robustly detected but there were no significant differences between Trim7 sufficient and deficient animals (**Figure 5A-5E**). These data demonstrate that removal of the STING pathway in mice does not reveal a role for Trim7 in mediating MNV infection in vivo.

**Figure 5.**
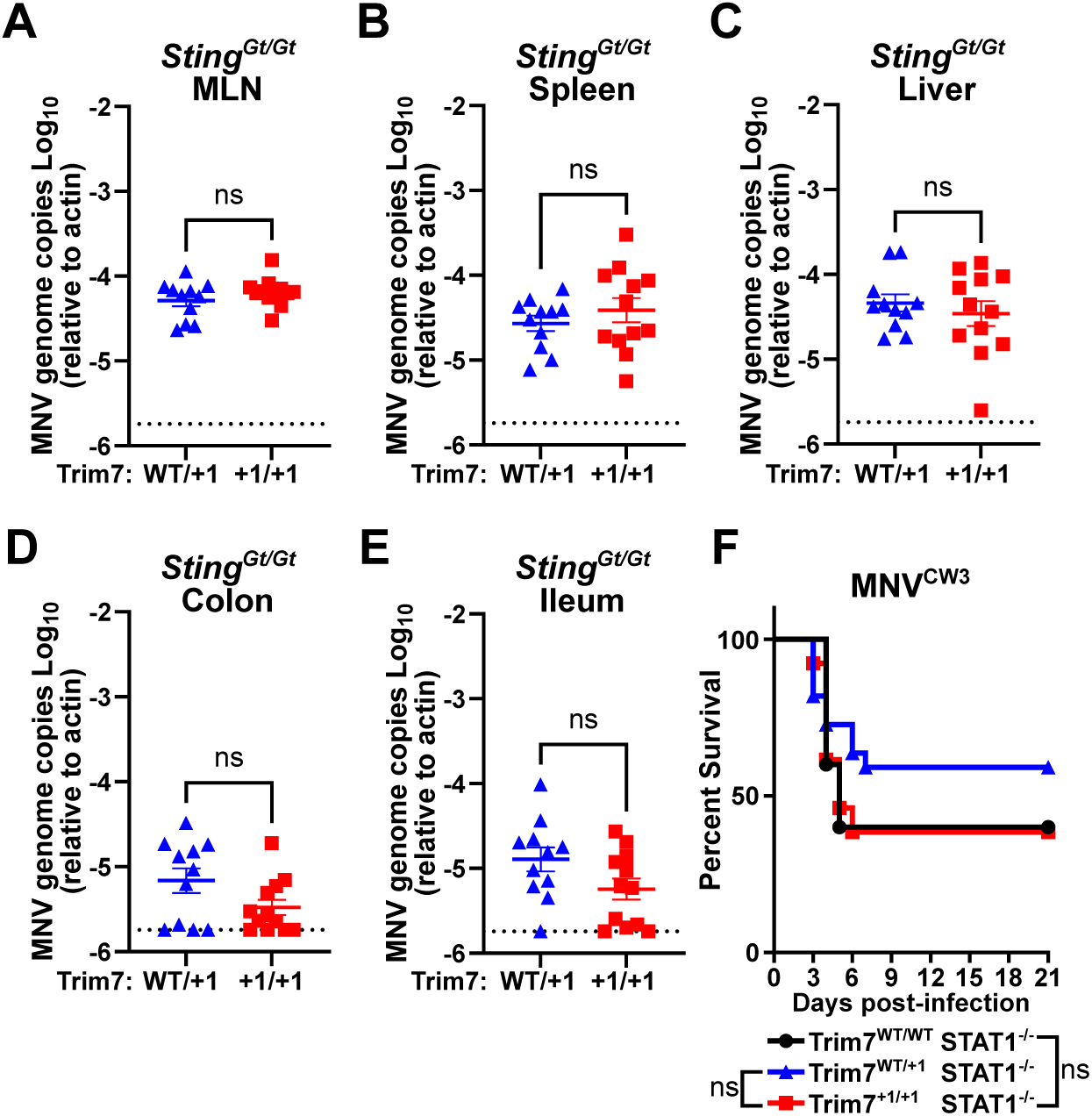
STING and STAT1 dependent innate pathways do not mask the role of Trim7 in restricting MNV^CW3^ infection. **A-E)** C57BL/6-Sting^Gt/Gt^ Trim7^+1/+1^ or littermate C57BL/6-Sting^Gt/Gt^ Trim7^WT/+1^ were inoculated with 5 × 10^6^ PFU of MNV^CW3^ and 7 days post-infection animals were sacrificed and MNV burden assed by measuring the genome copies in the MLN (**A**), spleen (**B**), liver (**C**), colon (**D**), and ileum (**E**) via qPCR. MNV genome levels were determined by qPCR and normalized relative to actin transcripts. Data are shown as mean ± S.D. from three independent experiments with 11-12 mice per group. ns, ** *P* < 0.01, *** *P* < 0.001, **** *P* < 0.0001, Mann-Whitney’s test. **F)** *STAT1^-/-^*/ Trim7^+1/+1^ and littermate control mice were inoculated with 1 × 10^3^ PFU of MNV^CW3^ and monitored daily for survival for 21 days post-infection (12-13 mice per group). Data from five independent experiments, analyzed using log-rank Mantel-Cox test.

Trim7 has also been reported to target MAVS, an RNA sensing pathway that is also responsible for regulating an innate immune response to MNV in vivo^20,41,45^. Both MAVS and STING converge upon type I and type III interferon (IFN) production which is a potent inhibitor of MNV replication both in vitro and in vivo^31^. Therefore, to eliminate the potential role of Trim7 regulating IFN induction, we crossed the Trim7^+1/+1^ mice onto a STAT1-/- background which eliminates all IFN signaling. STAT1 deficient mice are highly sensitive to MNV and succumb to infection unlike immunocompetent animals^12,28^. Trim7 sufficient and deficient animals lacking STAT1 were orally inoculated with a 1000 PFU of MNV^CW3^ and monitored for survival for 21 days. Deficiency of Trim7 did not affect the lethality caused by MNV^CW3^ in these mice (**Figure 5F**). Taken together our data finds a lack of a physiological role of Trim7 even in the absence of STING or STAT1.

## Discussion

Trim7 has emerged as a core component of host-virus interactions with both pro-viral and anti-viral activity based largely on compelling in vitro data. Due to the strong antiviral action of Trim7 overexpression in vitro, we hypothesized that Trim7 might have an important role in the clearance of MNV infection in vivo. However, our extensive studies found no significant role for Trim7 in MNV restriction using Trim7 deficient mice and different strains of MNV with distinct cellular and tissue tropisms. Trim7 deficiency did not have any effect on acute infection with MNV^CW3^ in systemic tissues, nor did it have any effect on persistent infection with MNV^CR6^ which primarily infects intestinal tuft cells.

MNV is efficiently controlled by the immune system and causes no symptomatic disease in immunocompetent adult mice. Both the innate and adaptive immune systems are necessary to clear infections^12^. Trim7 has been reported to influence host innate immune defenses due to its role in the degradation of STING and MAVS, leading to decreased inflammatory response to infection^20,21^. Thus, we tested the possibility that the physiological function of Trim7 is primarily focused on cytokine and innate immune responses rather than direct antiviral activity. In the context of acute MNV infection we did not observe any changes in the induction of pro-inflammatory cytokines, which contrasts with what others have reported for both DNA and RNA viruses^20,21^. However, neither STING nor MAVS have a C-terminal glutamine residue that has been structurally and biochemically demonstrated by multiple groups to be the hallmark of a Trim7 substrate^16–18^. Thus it is possible that neither STING nor MAVS are direct targets of Trim7, but the levels of these innate immune sensors are regulated indirectly by Trim7 in a context dependent manner. In addition to the lack of change in cytokine responses in infected mice lacking Trim7, we observe no impact of Trim7 on MNV infection patterns when STING or STAT1 are removed.

While this current study utilized multiple orthogonal approaches to probe for a physiological role for Trim7 in MNV infection, it does have several limitations. First, our data does not exclude the possibility that Trim7 may have a role in regulating MNV infections under very specific conditions that we did not test. Second, our data does not speak to whether other viruses whose protein products contain C-terminal glutamines are restricted by Trim7 in vivo. Lastly, our findings that the cytokine response to MNV infections is unaffected by Trim7, which is contrary to the models in the literature, may be a result of the specific virus we used or differences in the baseline physiology of our animals due to unidentified differences in animal facilities. Nevertheless, despite compelling evidence of the role of Trim7 in host defense in vitro, we see no physiological evidence for its activity in MNV clearance. Thus, caution should be exerted when classifying Trim7 as a broad acting antiviral recognizing the products of 3C-like protease cleavage events.

## Materials and Methods

### Mouse strains

All mouse experiments were conducted at University of Texas Southwestern Medical Center and approved by the University of Texas Southwestern Medical Center’s Institutional Animal Care and Use committees. C57BL/6J wild-type, C57BL/6J-Sting1gt/J and Stat1^−/−^ [B6.129S(Cg)-Stat1tm1Div/J]^38^ mice were originally purchased from Jackson Laboratories and bred in-housed under specific pathogen-free conditions, including devoid of murine norovirus. Two CRISPR KO lines for Trim7 deletion (BL6/Trim7^+1/+1^ and BL6ΔTrim) were used for our experiments. Trim7^+1/+1^ line was previously described^19^, and BL6ΔTrim7 line was made using the same strategy to create a larger deletion. Briefly, two sgRNAs targeting exon 1 (AGGACACGGATGGCGACTGT) and exon 7 (AGTTGACGCGGAAGGTGTAG) of the mouse Trim7 were generated, and co-microinjected with Cas9 mRNA into fertilized eggs of C57BL/6N mice^19^ (Figure 1A and 1B). Embryos were cultured overnight in M16 medium and were implanted into psuedopregnant foster mothers after reaching the 2-cell stage of development. Offspring were genotyped by PCR and sanger sequencing. Founder mice were bred with C57BL/6J mice to establish respective lines. All experiments were performed with gender-balanced littermate controls and independently replicated at least three times. Mice were used for infections between 6–10 weeks of age. Genotyping of the mice was done by real time PCR as described previously^19^.

### Western Blot

Indicated tissues from uninfected mice (Trim7^+1/+1^ and Trim7^Δ/Δ^) were collected, washed in ice-cold phosphate-buffered saline (PBS) and homogenized in RIPA lysis buffer (25mM Tris, 150mM NaCl, 1% IGEPAL, 0.5% sodium deoxycholate, 0.1% SDS) supplemented with Halt^TM^ protease inhibitor cocktail (Thermo Fisher Scientific) using a bead beater at 6800rpm for 1min. Homogenized samples were centrifuged at 13,000rpm for 20min at 4°C to remove cell debris. Protein concentrations of the supernatants were quantified using Coomassie Plus (Bradford) Assay Reagent (Thermo Fisher Scientific) normalized to a BSA standard curve on a BioTek Synergy LX Multimode Reader. Samples were then diluted and boiled with 2x Laemmli Sample Buffer (BioRad). For a positive control we used cell lysates overexpressing Trim7 isoforms. Lysates were resolved on SDS-PAGE gels and transferred to PVDF membranes.

Western blot was performed using the following antibodies: anti-Trim7 antibody produced in rabbit (Sigma SAB2106626; shown in **Figure 1B**), Trim7-antibody N-term (Abcepta AP11979a-ev), Trim7 polyclonal antibody (Bioss BS-9164R), Trim7 antibody C-term (GeneTex GTX24541), Anti-TRIM7 antibody produced in rabbit (Sigma HPA039213), Monoclonal Anti-GAPDH−Peroxidase antibody produced in mouse (Sigma G9295) and Anti-Mouse IgG (H+L), F(ab′)2 fragment peroxidase antibody in goat (Sigma SAB3701122).

### MNV infections in mice

MNV stocks were generated from plasmids encoding parental MNV^CW3^ (GenBank ID EF014462.1) or parental MNV^CR6^ (GenBank ID JQ237823) as described previously^39^. Viral stocks were obtained from plasmids expressing the complete genome of the viruses and purified and titered as previously described^40^. Genetic identity of viral stocks was confirmed by targeted sequencing.

Stocks of MNV^CW3^ P1 was diluted to 5×10^6^ PFU per 25uL of DMEM (with 5% fetal bovine serum) and inoculated per-orally in BL6/Trim7^+1/+1^, BL6/Trim7^Δ/Δ^, and BL6/Sting^Gt/Gt^ Trim7^+1/+1^ mice that were littermate-matched. Mice were euthanized at 7 days post-infection and mesenteric lymph nodes (MLN), spleen, liver, colon and ileum were harvested for RNA isolation. BL6/Stat1^-/-^Trim7^+1/+1^ mice and littermates were infected with MNV^CW3^ at 1000 PFU per animal, singly housed and monitored daily for survival for 21 days. MNV^CR6^ was inoculated in BL6/Trim7^+1/+1^ and littermate controls at 1×10^6^ PFU per animal. Mice were singly housed immediately after infection. Fecal samples were collected at days 3, 7, 14 and 21 post-infection. Mice were euthanized at 21 days post-infection and indicated tissues were harvested for RNA isolation.

### RNA extraction and qPCR assays

Quantification of MNV genomes from infected tissues was performed as previously described^40^. RNA was isolated from infected tissues using TRI Reagent (Sigma- Aldrich) with a Direct-zol kit (Zymo Research) following the manufacturers’ protocols. One µg of RNA was used for cDNA synthesis using a High-Capacity cDNA Reverse Transcription kit, following the manufacturer’s protocols (Thermo Fisher Scientific). TaqMan quantitative PCR (qPCR) for MNV was performed in triplicate on each sample and standard with forward primer 5’-GTGCGCAACACAGAGAAACG-3’, reverse primer 5’- CGGGCTGAGCTTCCTGC-3’, and probe 5’-6FAM-CTAGTGTCTCCTTTGGAGCACCTA-BHQ1-3’. TaqMan qPCR for Actin was performed in triplicate on each sample and standard with forward primer 5’- GATTACTGCTCTGGCTCCTAG-3’, reverse primer 5’-GACTCATCGTACTCCTGCTTG- 3’, and probe 5’-6FAM-CTGGCCTCACTGTCCACCTTCC-6TAMSp-3’.

RNA from infected fecal pellets was isolated using RNeasy Mini QIAcube Kit (Qiagen) and cDNA synthesis was performed using M-MLV Reverse Transcriptase kit (Invitrogen) using manufacturers’ protocols.

qPCR assays for cytokines and chemokines, and Trim7 were designed from IDT and the assay was performed in triplicate on each sample and standards with forward primer, reverse primer and probes as listed below:

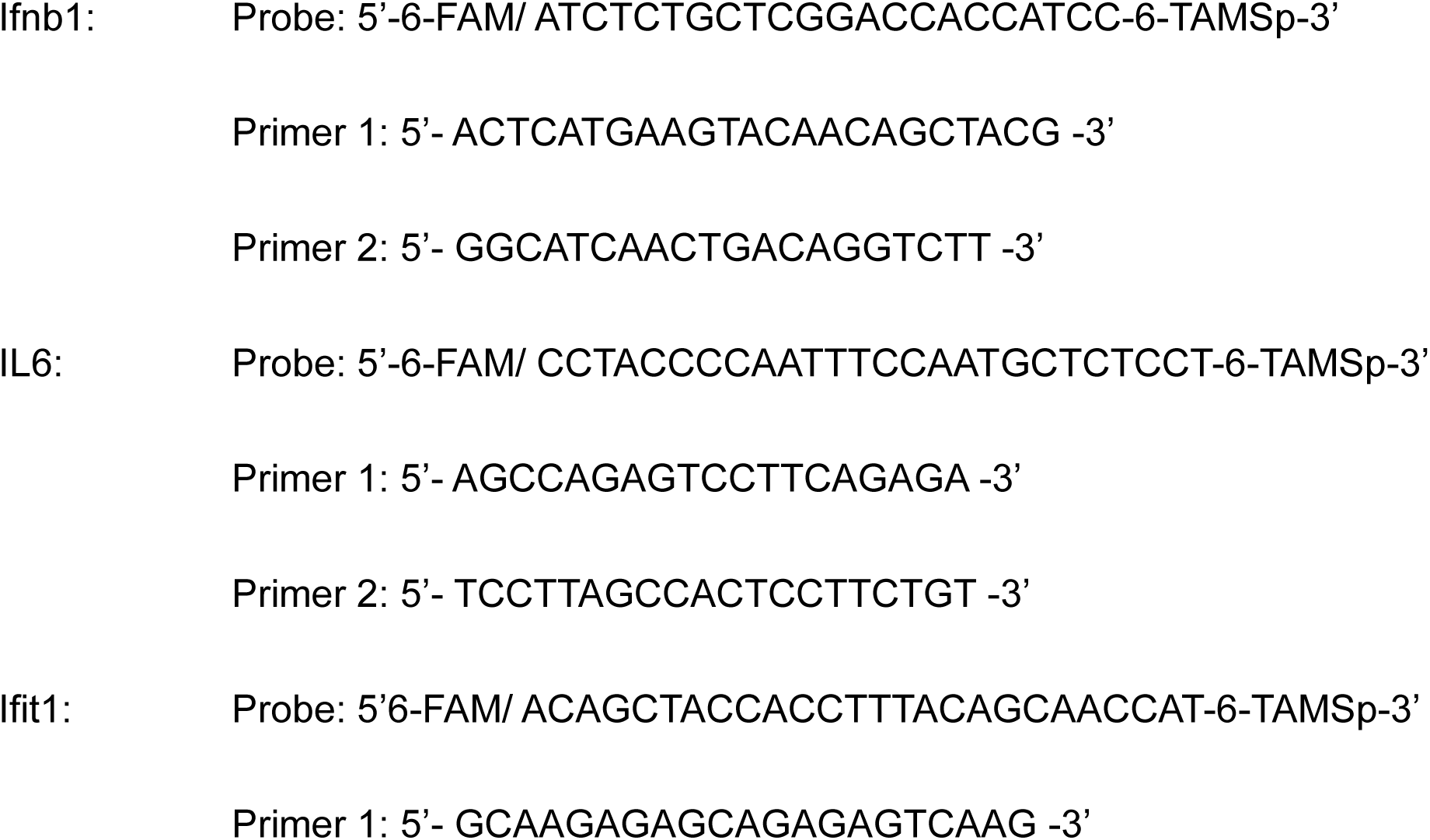
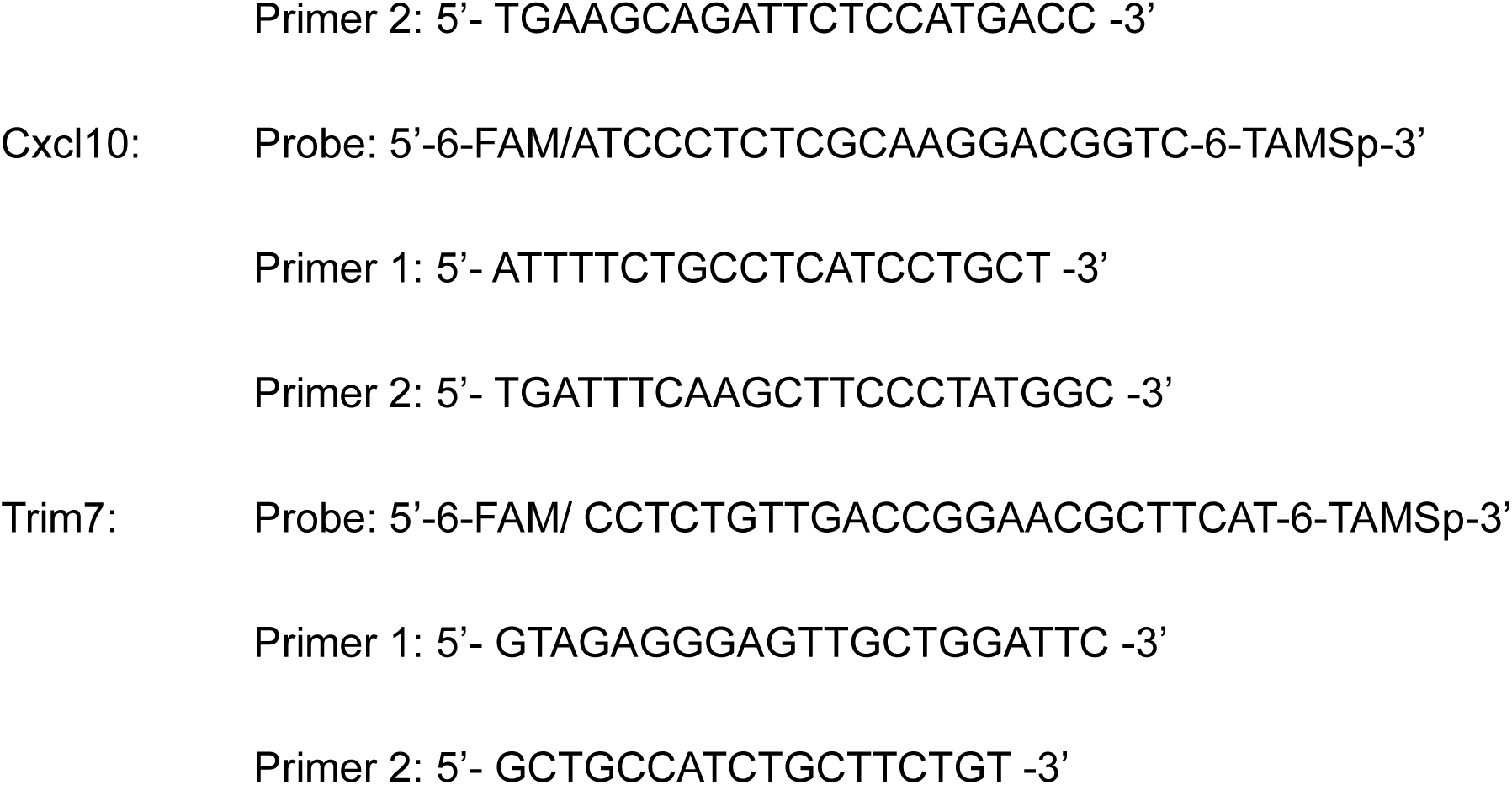

These transcript levels were normalized to actin and indicated as fold change relative to infected BL6/WT tissues.

## Acknowledgements

We would like to thank John Schoggins and all members of the Orchard Lab for helpful discussions. This work was supported by NIH grant 5R01DK133231 (R.C.O.). The work was supported in part by the Division of Intramural Research, National Institutes of Health, National Institute of Allergy and Infectious Diseases.

## Author Contributions

M.A.S. designed the project, performed experiments and helped draft the paper. L.R.P., M.C.O. S.J.R., G.L.S. and S.M.B helped perform experiments and provided critical reagents. R.C.O. conceptualized the project, provided supervision, and helped write the paper. All authors read and edited the manuscript.

## Disclosures

The authors have no financial disclosures.

